# Is it impossible to acquire absolute pitch in adulthood?

**DOI:** 10.1101/355933

**Authors:** Yetta Kwailing Wong, Kelvin F. H. Lui, Ken H.M. Yip, Alan C.-N. Wong

## Abstract

Absolute pitch (AP) refers to the rare ability to name the pitch of a tone without external reference. It is widely believed that AP is only for the selected few with rare genetic makeup and early musical training during the critical period. Accordingly, acquiring AP in adulthood is impossible. Previous studies have not offered a strong test of the effect of training because of issues like small sample size and insufficient training. In three experiments, adults learned to name pitches in a computerized and personalized protocol for 12 to 40 hours. They improved considerably, with a continuous distribution of learning progress among them. 14% of the participants (6 out of 43) were able to name twelve pitches at accuracy of 90% or above, comparable to that of ‘AP possessors’ as defined in the literature. In general, AP learning showed classic characteristics of perceptual learning, including generalization of learning dependent on the training stimuli, and sustained improvement for at least one to three months. Overall, the finding that AP continues to be learnable in adulthood calls for reconsidering the role of learning in the occurrence of AP. The finding also pushes the field to pinpoint and explain, if any, the differences between the aspects of AP more trainable in adulthood and the aspects of AP that are potentially exclusive for the few exceptional AP possessors.

---

Absolute pitch (AP) refers to the ability to name the pitch of a tone (e.g., naming a tone as “C”) or to produce it without external reference tones (Takeuchi & Hulse, 1993; W. D. Ward, 1999). While the majority of us can effortlessly identify a countless number of faces, objects, and visual and auditory words, most people find it very difficult to name the twelve pitches, and professional musicians are of no exception (Athos et al., 2007; Levitin & Rogers, 2005; Zatorre, 2003). The most extreme estimate states that, in every 10000 people, there is one ‘AP possessor’ who can perform AP judgment accurately and effortlessly (Takeuchi & Hulse, 1993). This rare ability is considered a special talent and endowment for gifted musicians (Deutsch, 2002; Takeuchi & Hulse, 1993; W. D. Ward, 1999). The genesis of AP has therefore been a perplexing research topic among musicians, psychologists and neuroscientists for more than a century (Deutsch, 2002; Levitin & Rogers, 2005; Takeuchi & Hulse, 1993; W. D. Ward, 1999).

At the cognitive level, a widely accepted hypothesis suggests that AP involves two stages of processing (Levitin, 1994; Levitin & Rogers, 2005). The first stage is the experience-based AP memory that general listeners can also establish (Halpern, 1989; Levitin, 1994; Schellenberg & Trehub, 2003). The second stage is the ability to associate the represented AP memory with verbal labels, which is somehow mastered by the ‘AP possessors’ only (Brancucci, Dipinto, Mosesso, & Tommasi, 2009; Deutsch, 2002; Levitin & Rogers, 2005; Schellenberg & Trehub, 2003; Vanzella & Schellenberg, 2010). In other words, the bottleneck of AP performance is in assigning verbal labels to the tones rather than establishing AP memory representation per se.

At the neural level, AP has been associated with distinct neural markers. In functional neural imaging, ‘AP possessors’ have shown an increased activity in the superior temporal gyrus (Ohnishi et al., 2001; Schulze, Gaab, & Schlaug, 2009; Wengenroth et al., 2013; S. J. Wilson, Lusher, Wan, Dudgeon, & Reutens, 2009) and the left dorsal lateral prefrontal cortex (Bermudez & Zatorre, 2005; Zatorre, Perry, Beckett, Westbury, & Evans, 1998). AP has also been linked with various event-related potentials (ERPs), including N1 (Itoh, Suwazono, Arao, Miyazaki, & Nakada, 2005; Pantev et al., 1998; Wu, Kirk, Hamm, & Lim, 2008), P2a (Wengenroth et al., 2013), and P3 (Hantz, Crummer, Wayman, Walton, & Frisina, 1992; Hirose et al., 2002; Klein, Coles, & Donchin, 1984; Rogenmoser, Elmer, & Jäncke, 2015). Other structural and functional connectivity differences have also been identified, including the size asymmetry of the planum temporale (Keenan, Thangaraj, Halpern, & Schlaug, 2001; Schlaug, Jancke, Huang, & Steinmetz, 1995), the architecture of the superior longitudinal fasciculus (Oechslin, Imfeld, Loenneker, Meyer, & Jäncke, 2009), and functional connectivity of different brain regions (Jäncke, Langer, & Hänggi, 2012; Loui, Li, Hohmann, & Schlaug, 2011; Loui, Zamm, & Schlaug, 2012).

## The role of genes and the critical period

How can we explain the development of AP in the ‘AP possessors’? One influential theory suggests that AP develops if two prerequisites are fulfilled (Bachem, 1940; Baharloo, Johnston, Service, Gitschier, & Freimer, 1998; Chin, 2003; Drayna, 2007; Levitin & Rogers, 2005; Ross, Olson, Marks, & Gore, 2004; Takeuchi & Hulse, 1993; Trainor, 2005; Zatorre, 2003). The first prerequisite is the rare genetic disposition to AP. It is reported that AP runs in families, as siblings of ‘AP possessors’ are also more likely to be ‘AP possessors’ (about 14 - 48% of the ‘AP possessors’ reported to have a sibling or first-degree relatives that was also an ‘AP possessor’; Baharloo et al., 1998; Baharloo, Service, Risch, Gitschier, & Freimer, 2000; Gregersen, Kowalsky, Kohn, & Marvin, 1999, 2001). AP is often considered a relatively ‘clean’ cognitive phenotype (Baharloo et al., 2000; Gregersen et al., 2001), and some proposed that AP may be subserved by a single gene (Drayna, 2007).

The second prerequisite is an early onset of musical training that is within the critical period in childhood. A strong definition of critical period refers to the early period of life during which experience is essential for normal development and leads to permanent changes in brain functions and behavior (Knudsen, 2004)^1^, and we based the current discussion on this definition because this description fits well with the widespread belief of AP (Bachem, 1940; Baharloo et al., 1998; Chin, 2003; Drayna, 2007; Levitin & Rogers, 2005; Ross et al., 2004; Takeuchi & Hulse, 1993; Trainor, 2005; Zatorre, 2003). The critical period of AP is thought to be similar to that of language development (Chin, 2003; Deutsch, Dooley, Henthorn, & Head, 2009). Supporting evidence comes from the large-scale survey studies that the average onset of musical training of ‘AP possessors’ was 5.4 years, which was about 2.5 years earlier than that of ‘non-AP possessors’ (Gregersen et al., 1999; see also Baharloo et al., 1998; Gregersen, Kowalsky, Kohn, & Marvin, 2001). Importantly, AP training was considered relatively successful in young children (Crozier, 1997; Miyazaki & Ogawa, 2006; Sakakibara, 2014). In contrast, while AP can improve to some extent in adulthood with deliberate practice (Brady, 1970; Cuddy, 1968, 1970; Hartman, 1954; Meyer, 1899; Mull, 1925; Russo, Windell, & Cuddy, 2003; Van Hedger, Heald, Koch, & Nusbaum, 2015; Wedell, 1934), there is no convincing evidence that adults can attain a performance level comparable to the AP possessors through training (Bachem, 1940; Levitin & Rogers, 2005; W. D. Ward, 1999). According to this influential theory, most professional musicians fail to acquire AP because they fail to start their music training within the critical period and/or because they do not carry the specific genes. Training AP in adulthood, when the critical period of acquiring AP has long passed, should be practically impossible (Bachem, 1940; Crozier, 1997; Trainor, 2005; but see Gervain et al., 2013 for the possibility of reopening the critical period by taking a drug).

The concept of the critical period well explains the development of basic functions and structure of different sensory systems. For example, in vision, it is well-known that depriving the visual input into one eye led to atrophy and reduced neural activity in lateral geniculate body and reduced ocular dominance in the visual cortex of young cats, while similar results were not observed in deprived adult cats (Hubel & Wiesel, 1970; Wiesel & Hubel, 1963). In hearing, disrupting the normal pattern of sound input during the early life of rats led to abnormal development of the tonotopic maps and functions of the auditory cortex, while similar exposure did not cause significant changes in deprived older rats (Zhang, Bao, & Merzenich, 2002). In touch, depriving tactile inputs after birth led to abnormalities in the functions of the somatosensory cortex (Simons & Land, 1987). Overall, normal development of structure and functions of the sensory cortices is fundamentally determined by the environmental inputs and experience during early period of life (but see Hooks & Chen, 2007 for recent challenges of this view).

However, evidence is less as clear for high-level cognitive abilities. In vision, different types of higher-level visual abilities, such as acuity, stereopsis and crowding, continue to develop in adulthood (Daw, 1998). In musical development, various degrees of plasticity in adulthood has been observed during the acquisition of musical pitch structure (e.g., consonance and dissonance, scale structure and harmonics) and the structure and functions of the auditory and motor cortex relevant to musical training, and there was no direct evidence for critical periods for these abilities (see review in Trainor, 2005). In language acquisition, it has been argued that there is a critical period for language development, especially in terms of the limitations for late learners to attain a native-like accent (Patkowski, 1990; Scovel, 1988). However, subsequent research demonstrated that it is possible for later learners, who started to acquire a second language after twelve years old, to attain native accents in various languages, including English, French and Dutch (Flege, Munro, & MacKay, 1995; see review in Singleton, 2001). Another example comes from synesthesia, in which the perception of a stimulus (e.g., a black letter) consistently evokes experience that is not physically present (e.g., seeing the color ‘red’ on a black letter; J. Ward, 2013). It is hypothesized that all infants are born to be synesthetes, and this condition persists into adulthood for some individuals because of reduced pruning during early childhood (Maurer & Mondloch, 2006). Synesthesia tends to aggregate in families (J. Ward & Simner, 2005), and has a close genetic link with AP (Gregersen et al., 2013). Interestingly, recent work suggests that adults can be trained to acquire synesthetic experiences (Bor, Rothen, Schwartzman, Clayton, & Seth, 2015). This is yet another example of complex human behavior, previously thought to be constrained by genes and early development, that might in fact be learnable during adulthood. Intriguingly, AP appears to present a particularly strong case for the theoretical proposal of the critical period constraining the development of high-level cognitive abilities, in contrast to other examples of high-level cognitive abilities (Zeanah, Gunnar, McCall, Kreppner, & Fox, 2011).

A closer look into the literature of AP indicates that direct evidence for a critical period constraining AP development is weak. Similar to accent acquisition, some cases of AP were identified with later onset of musical training in the normal population (about 3-4% in the age groups of 9-12 and beyond 12; Baharloo et al., 1998) and in the individuals with the Williams Syndrome (3 out of 5 ‘AP possessors’ with the Williams Syndrome started musical training at 8 years old or later; Lenhoff, Perales, & Hickok, 2001). These suggest that early onset of musical training during the critical period is not essential for developing AP (Gregersen et al., 2001). If the critical period is not essential and the genetic contribution of AP is only moderate, these two factors are unlikely to be the whole picture that describes AP development (Baharloo et al., 1998, 2000, Gregersen et al., 1999, 2001). What other factors could explain the genesis of AP?

### The role of experience

A possible alternative is that AP is developed through perceptual learning. Perceptual learning refers to the long-term improvement on a perceptual task as a result of perceptual experience (Fahle & Poggio, 2002; Goldstone, 1998; Sasaki, Nanez, & Watanabe, 2010). It has been repeatedly demonstrated that humans extract information from environmental inputs and fine-tune their perceptual representations accordingly in different sensory modalities, including the visual (Fiorentini & Berardi, 1980; Karni & Sagi, 1993), auditory (Fujioka, Ross, Kakigi, Pantev, & Trainor, 2006; Kraus & Banai, 2007), somatosensory (Sathian & Zangaladze, 1997) and olfactory domains (D. A. Wilson & Stevenson, 2003). Perceptual learning occurs even when the stimuli are irrelevant to the task on hand (Seitz & Watanabe, 2009), for the non-diagnostic and task-irrelevant information for categorizing trained stimuli (Y. K. Wong, Folstein, & Gauthier, 2011), or when participants are unaware of the information carried by the stimuli (Tsushima, Sasaki, & Watanabe, 2006; Watanabe, Nanez, & Sasaki, 2001). These demonstrate how the perceptual system is constantly tuned to environmental inputs. Consistent with the perceptual learning hypothesis of AP, it is well-established that AP is shaped by experience (Takeuchi & Hulse, 1993; Y. K. Wong & Wong, 2014). For example, better pitch naming is often observed in musicians when the testing conditions better match one’s prior experience, such as using tones in the timbre of one’s own instrument (see review in Takeuchi & Hulse, 1993), tones in a highly used pitch like ‘A4’, which is the standard tuning pitch in orchestras (Levitin & Rogers, 2005; Takeuchi & Hulse, 1993), tones presented in a multisensory testing context similar to one’s musical training (Y. K. Wong & Wong, 2014), and tones associated with the more frequently used white keys than black keys (Athos et al., 2007; Miyazaki, 1989, 1990; Takeuchi & Hulse, 1993). Even performance in ‘AP possessors’ can be disrupted with recent listening experience with detuned music (Hedger, Heald, & Nusbaum, 2013). Experience also explains why non-musicians can recognize the starting tones of songs or melodies that are highly familiar in an absolute manner, which indicates that non-musicians forms AP memory of tunes based on repeated exposure (Levitin, 1994; Schellenberg & Trehub, 2003).

Previous training studies might have failed to convince researchers that AP is trainable in adulthood because of various issues. For example, the training duration was very short in some studies (roughly 1-4 hours; Cuddy, 1970; Mull, 1925; Van Hedger et al., 2015), which might have limited the potential of observing larger effects in AP learning. Some training studies used a very small sample size (3 or less participants per condition; Brady, 1970; Cuddy, 1968; Hartman, 1954; Meyer, 1899; Mull, 1925; Wedell, 1934) and involved self-training of the authors of the manuscripts (Brady, 1970; Meyer, 1899). Other training studies were difficult to interpret, e.g, with only a binary judgment on a single tone learned (e.g., “C” or not “C”; Mull, 1925; Russo et al., 2003), or with insufficient information provided for interpreting the performance of the participants (Lundin, 1963). Therefore, despite the apparently substantial AP improvement attained in some of these studies (e.g., Brady, 1970; Lundin, 1963), it remains unclear whether AP can be acquired in adulthood. Furthermore, previous training studies might have lacked key components of effective training regimes, such as computerized stimulus presentation and feedback protocols that enable indiviudalized training progress, incentive for sustaining motivation of the training, etc, which might have collectively limited the potential AP improvement during training.

In this study, we tested whether AP acquisition in adulthood, i.e., attaining a performance level comparable to that of real-world ‘AP possessors’, is possible through perceptual training, and whether the training-induced improvements are consistent with that of perceptual learning. We defined AP to be the ability to name pitches accurately without external references, which is one of the most common definitions of AP in the literature (Levitin & Rogers, 2005; Takeuchi & Hulse, 1993; W. D. Ward, 1999). To test whether AP can be acquired, we first needed to define the performance level of real-world ‘AP possessors’. However, our survey of the literature revealed that the definition of ‘AP possessors’ was variable and arbitrary in previous work (e.g., using self-report or different performance-based measurements, with different scoring methods and cut-off points; see Methods). In this study, we arbitrarily adopted a relatively stringent training criterion of AP - being able to name, without any reference, all of the twelve pitches that constitute an octave at 90% accuracy, with semitone errors considered incorrect. Our survey of the literature indicates that participants who have passed our training would also be considered ‘AP possessors’ in most of the published AP studies that adopted performance-based definition of ‘AP possessors’ (see Methods of Experiment 1). This indicates that individuals who have passed our training have acquired AP performance comparable to that of ‘AP possessors’ as defined in the literature.

To address our question, we designed different training regimes of AP by optimizing various factors, including the use of extended training duration, larger sample sizes, modern computerized protocols and measures to provide individualized training and to sustain motivation, e.g., by gamifying the training and including monetary reward for the training progress. In three experiments, we trained 43 adults to name the pitch of tones with different combinations of timbres and octaves for 12-40 hours in laboratory and mobile online settings (Table 1). If specific genetic disposition and an early onset of music training are essential for AP acquisition, AP training in adulthood should be largely in vain resulting in very limited improvement in all participants. Alternatively, if AP can be trained in adulthood as a type of perceptual learning, it should be possible, at least for some individuals, to attain a performance level similar to that of real-world ‘AP possessors’.

In addition, if AP is developed through perceptual learning, then AP learning should demonstrate several characteristics that match well with that of perceptual learning studies. First, perceptual learning typically leads to performance enhancement after training (Fahle & Poggio, 2002; Goldstone, 1998). In this study, we examined whether AP performance improved by measuring the number of pitches that participants learned to name accurately without feedback through training (see Methods). Second, we examined whether the degree of generalization of AP learning changed as a function of the octaves and timbre involved in training. Although perceptual learning is often regarded as highly specific to the training stimuli (Fahle & Poggio, 2002; Watanabe & Sasaki, 2015), especially when the training is difficult and supported by more specific inputs (Ahissar & Hochstein, 2004), generalization of learning is often present in various untrained conditions to different degrees (Banai & Lavner, 2014; Fahle & Poggio, 2002; Watanabe & Sasaki, 2015; Wright, Buonomano, Mahncke, & Merzenich, 1997). Instead of simply regarding perceptual learning as ‘specific’, the degree of learning specificity and generalization during perceptual training is best conceptualized by understanding the psychological space involved in training and testing – whether the testing space overlaps with the training space would affect the extent of generalization (Y. K. Wong et al., 2011; see also Nosofsky, 1986, 1987; Palmeri & Gauthier, 2004). Consistently, we predicted that including training tones in more octaves and timbres (as in Experiment 1 and 3) should cover larger regions of the psychological space, and thus lead to better generalization to untrained octaves and timbres, while including training tones only in a single octave and timbre (as in Experiment 2) should result in more specific AP learning. Third, perceptual enhancement in perceptual learning is relatively long-lasting, often in the time scale of months or even years (Fahle & Poggio, 2002; Goldstone, 1998; Karni & Sagi, 1993; Watanabe & Sasaki, 2015). In three experiments, we tested whether any AP improvement sustained for one to three months after the end of training. Finally, we examined whether musicians, who have been frequently exposed to musical tones during musical training, would learn AP more efficiently than non-musicians (Experiment 2). To preview the results, all participants improved, with musicians learning to name more pitches than non-musicians. While there is a continuous distribution of learning progress among them, 14% of the participants (6 out of 43) attained a performance level comparable to that of ‘AP possessors’ typically identified in the literature. Learning generalized to the untrained octaves and timbre more when the training tones involved more octaves and timbres (in Experiments 1 & 3), and was more specific when the training tones involved only a single octave and timbre (Experiment 2), and sustained AP improvement for at least one to three months after training.

**Table 1.** Details of the training protocols in the three experiments. N indicates the number of participants in each experiment. ‘Mus/Non-Mus’ indicates the number of musicians and non-musicians in each experiment respectively.

| <u>Experiment</u> | <u>Participants</u> |  | <u>Training</u> |  |  |  | <u>Test for Generalization</u> |  |
| --- | --- | --- | --- | --- | --- | --- | --- | --- |
|  | <u>N</u> | <u>Mus/Non-Mus</u> | <u>Mode</u> | <u>Duration(Hr)</u> | <u>Octave</u> | <u>Timbre</u> | <u>Octave</u> | <u>Timbre</u> |
| 1 | 10 | 7/3 | Lab | 12 | 4,5 | synthetic tones,<br>piano | 3,6 | violin |
| 2 | 22 | 10/12 | Lab | 15 | 4 | synthetic tones | 5 | piano,<br>violin |
| 3 | 11 | 6/5 | Online | 40 | 4,5 | piano,<br>violin,<br>clarinet | 6 | synthetic |

## Experiment 1

In Experiment 1, we used a more optimized regime of AP training compared with previous studies (Brady, 1970; Cuddy, 1968, 1970; Meyer, 1899; Van Hedger et al., 2015), with extended training duration, larger sample sizes, modern computerized protocols and measures to sustain motivation. The aim was to test whether learning AP in adulthood is possible, and whether the training-induced improvement follows the characteristics of perceptual learning. Since how much training contributes to a sufficiently rigorous training to enable AP acquisition in adulthood is an untested empirical question, we decided to conduct a 12-hour training in this experiment, which was comparable to some previous perceptual learning studies in our laboratory (A. C. Wong, Palmeri, & Gauthier, 2009; Y. K. Wong et al., 2011).

## Methods

### Participants

Ten adults were recruited at City University of Hong Kong and completed the training. They included 2 males and 8 females, who were 23.1 years old on average (*SD* = 4.50). Seven of them were trained in music for 2-10 years, with the major instrument being piano (*N* = 5), violin (*N* = 1) and flute (*N* = 1). Three were non-musicians who were not formally trained with music before. One additional participant dropped out in the middle of the training and was excluded from all analyses. All participants filled out a questionnaire about their musical training background, including the musical instruments and the highest ABRSM exam passed, and reported if they regarded themselves as ‘AP possessors’. They received monetary compensation for the training and testing. Informed consent was obtained according to the Ethics Committee of City University of Hong Kong.

The sample size was estimated based on a recent AP training study (Van Hedger et al., 2015) using GPower 3.1.9.2. In this study, a large effect size was observed for the training improvement in adults (pretest vs. posttest; *f* = 1.34). Using the same *f*, the sample size required to detect any training effect at *p* = .05 with a power of 0.95 was 5 participants. To be more conservative, we recruited 10 participants. This sample size was also consistent with that used in previous perceptual training studies (Chung & Truong, 2013; Y. K. Wong et al., 2011).

### Materials

The experiment was conducted on personal computers using Matlab (Natick, MA) with the PsychToolbox extension (Brainard, 1997; Pelli, 1997) at the Cognition and Neuroscience Laboratory at City University of Hong Kong. Participants were requested to bring their own earphone to the training and testing. They adjusted the volume to a comfortable level before the training or testing started.

In Experiment 1, 120 tones from octaves three to six were used. They were synthetic tones and piano tones in octaves three to six, and violin tones in octaves four to five. The synthetic tones were identical to those in prior AP tests, and was generated by summing a series of sinusoidal waveforms including the fundamental frequency and harmonics (Bermudez, Lerch, Evans, & Zatorre, 2009). The piano tones were recorded with an electric keyboard (Yamaha S31). The violin tones were recorded by a volunteer violinist in a soundproof room. The precision of the tones was checked during recording by a tuner. The sound clips were 32-bit with a sampling rate of 44100Hz. They were edited in Audacity such that they lasted for 1 second with a 0.1-second linear onset and 0.1-second linear offset and were matched with similar perceptual magnitude.

### Absolute pitch training

The training included 48 tones from two octaves (C4 to B5) in two timbres of synthetic and piano tones. A pitch-naming task was used. During each trial, an isolated tone was presented for 1s. Then, an image showing the pitch-to-key mapping was presented (Figure 1A). Participants were required to name the pitch of the presented tone by key press within 5s.

**Figure 1.**
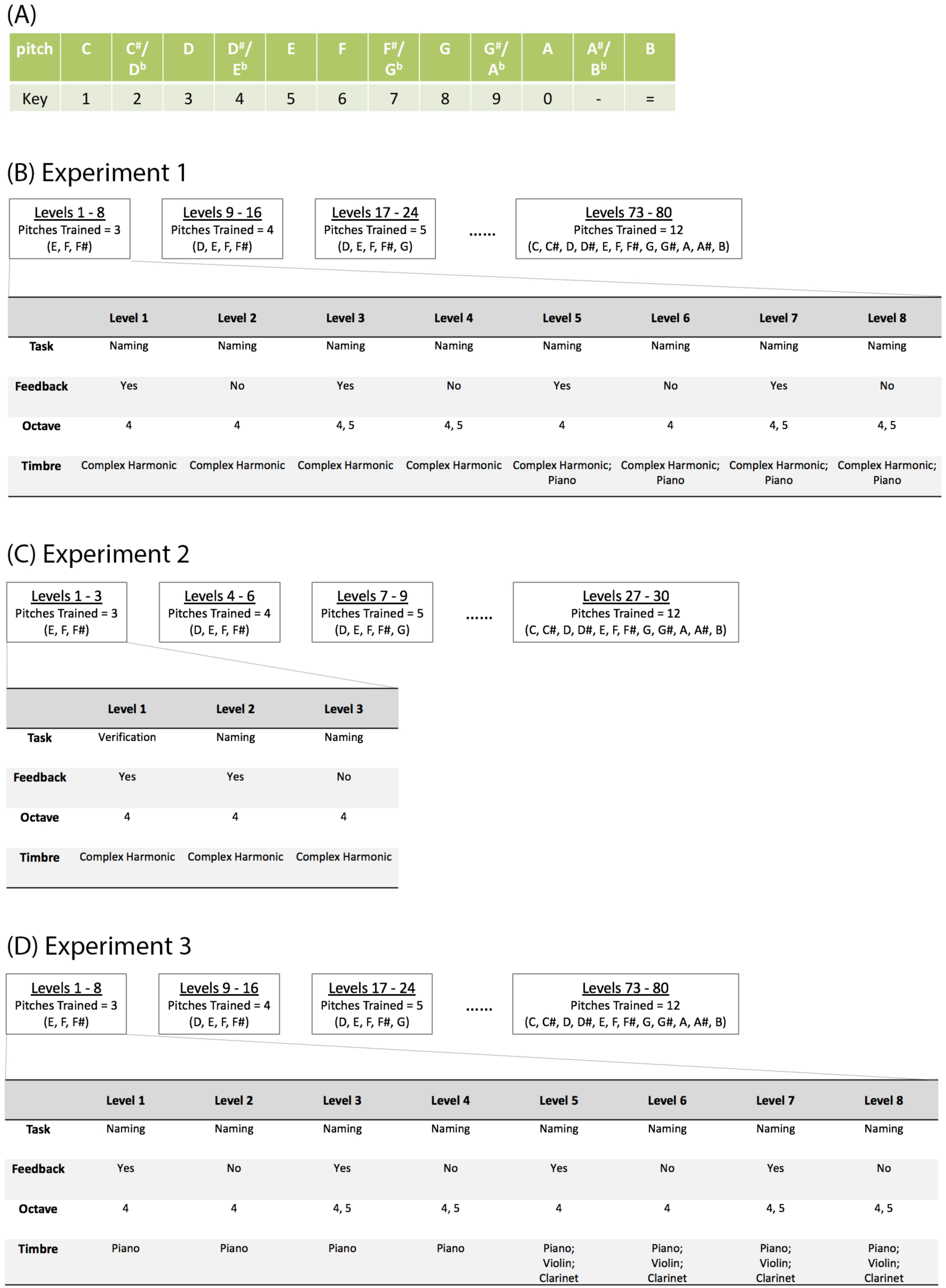
The design of the AP training in the three experiments. (A) shows the image probing for pitch naming responses by key press during the training and the test for generalization. During the training, only the pitch-to-key mapping for the trained tones was included in the image. During the test for generalization, the image showing the mapping for all twelve pitches was always used. (B-D) show the design of the training levels in terms of the task, presence of trial-by-trial feedback, and the octave and timbre of the training tones of Experiment 1, 2 and 3 respectively.

The training was gamified and structured with different levels. If participants achieved 90% accuracy for a certain level, they would proceed to the next level; otherwise they would stay at the same level. The training was completed by finishing 12 hours of training or by passing all 80 levels with 90% accuracy. Participants finished one hour of training per day. They were trained on at least four days per week and finished the training in three weeks.

The 80-level training protocol was organized into ten 8-level parts with an increasing number of pitches (from 3 pitches in the first 8 levels to 12 pitches in the last 8 levels; Figure 1B). Each eight-level part consisted of four types of levels, which included tones that were progressively richer in timbres and octaves. Each of the four types of levels was repeated twice, once with trial-by-trial feedback provided, and then once without feedback. For example, participants began the training with three pitches (E, F and F#). At levels 1-2, synthetic tones in these three pitches in octave 4 were included, with feedback provided at level 1 and then without feedback at level 2. At levels 3-4, synthetic tones in both octaves 4 and 5 were included with feedback and then without feedback. At levels 5-6, synthetic tones and piano tones in octave 4 were included with feedback and then without feedback. At levels 7-8, synthetic tones and piano tones in octaves 4 and 5 were included with feedback and then without feedback. At the no-feedback levels, participants were not provided with any feedback of the correctness of the tones. These no-feedback levels served as mini milestones for participants’ AP performance at 90% accuracy. If they achieved 90% accuracy at the 8^th^ level, a new pitch was added into the training set, with which they went through the same 8-level part again. Each level included 20 trials, with tones distributed as evenly as possible among the training pitches, octaves and timbres. Semitone errors were considered errors in the training. Before each level, participants were allowed to freely listen to sample tones of the training pitches as many times as they preferred before proceeding to the training. Each training session lasted for an hour, in which individual participants might have finished different numbers of training trials depending on their pace of learning (e.g., the amount of time spent on the training trials or on sample tone listening).

Normally participants earned one point for each correct answer in each trial. To motivate participants, a special trial that was worth three points randomly appeared with a chance of 1/80. Also, participants were given 1, 2 and 3 tokens if they achieved 60%, 75% and 90% accuracy at a training level respectively. With ten tokens, participants would obtain a chance to initiate the three-point special trial when preferred. At most three chances of initiating these special trials at one level were allowed. The special trials did not appear and could not be initiated during the no-feedback levels. This ensured that participants performed the no-feedback levels without any scoring assistance.

### Definition of AP

In order to compare the trained AP performance with that of real-world ‘AP possessors’, we surveyed the literature of AP on Web of Science on 19^th^ April, 2017 with the term ‘absolute pitch’ in the topic and identified 110 empirical papers. Unfortunately, there was not a single objective definition of the performance level of ‘AP possessors’. Instead, these papers used highly varied definitions of ‘AP possessors’, including self-report, AP performance measurements, or relative performance on AP tasks between participants (such as 3 SDs higher in AP accuracy than ‘non-AP possessors’). We focused on the 66 publications that defined AP objectively based on AP performance instead of self-report, and found that these papers adopted highly varied performance measures to define ‘AP possessors’, including cut-off points and scoring methods (taking semitone errors as correct, partially correct or incorrect; using accuracy or the average size of errors, etc.). Given this variability in definition, we did not see any strong reasons to adopt any single definition of ‘AP possessors’ based on some particular publications. In this study, we arbitrarily adopted a relatively stringent training criterion of AP, i.e., having achieved 90% pitch naming accuracy with all of the 12 pitches and with semitone errors considered incorrect. We asked whether participants who have passed our training would be considered ‘AP possessors’ according to the definition specified in each of the 66 papers. We recalculated the participants’ performance if needed. We observed that our successfully trained participants would be considered ‘AP possessors’ in 83.3% (55 out of 66) of those papers using objective definitions, meaning that their AP performance was representative of and comparable to that of ‘AP possessors’ as defined in the literature.

### Test for generalization

The test for generalization was performed before and within three days after training to examine how well the pitch-naming abilities generalized to untrained octaves and timbres. 120 tones in octaves 3 to 6 were used, in which octaves 4 and 5 were trained, and 3 and 6 untrained. Three timbres were included, with synthetic and piano tones as trained timbres, and violin tones as an untrained timbre. The tones were presented in three conditions, either with trained octave and timbre, trained octave and untrained timbre, or untrained octave and trained timbre. During each trial, a tone was presented for 1s. Then an image that mapped the 12 pitch names to 12 keys of the keyboard, which was the same as that used in the training, was presented (Figure 1A). Participants were required to name the pitch of the presented tone by key press within 5s. Each tone was presented twice, leading to 240 trials in total. The trials were presented in randomized order. No practice trials were provided in these tests. The dependent measure was the precision of pitch naming, i.e., the average error in semitone of participants’ responses relative to the correct responses. A trial without a response will be assigned an error of 3 semitones (the expected error one would have by complete guessing). We used the average error instead of the general naming accuracy because measuring the size of the judgment errors additionally informs the precision of pitch naming performance of the individuals, which is more informative than the binary correctness of the responses as measured by general naming accuracy. An identical test was performed a month later to test whether the AP learning sustained for at least a month.

## Results

### Learning of AP

In general, participants made substantial progress in learning to name pitches. At the end of training, they were able to name on average 8.1 pitches (out of 12; range = 5 to 12; SD = 2.28) at 90% accuracy without any externally provided reference tones under the stringent scoring criterion of taking semitone errors as incorrect (Figure 2A). Importantly, one of the ten participants passed all levels of training, meaning that he was able to name all of the twelve pitches at 90% accuracy without any externally provided reference tones. With this level of AP performance, he would be recognized as an ‘AP possessor’ in most of the empirical papers that adopted an objective performance-based definition of AP (see Methods). These suggest that he has acquired AP through perceptual learning in adulthood.

**Figure 2.**
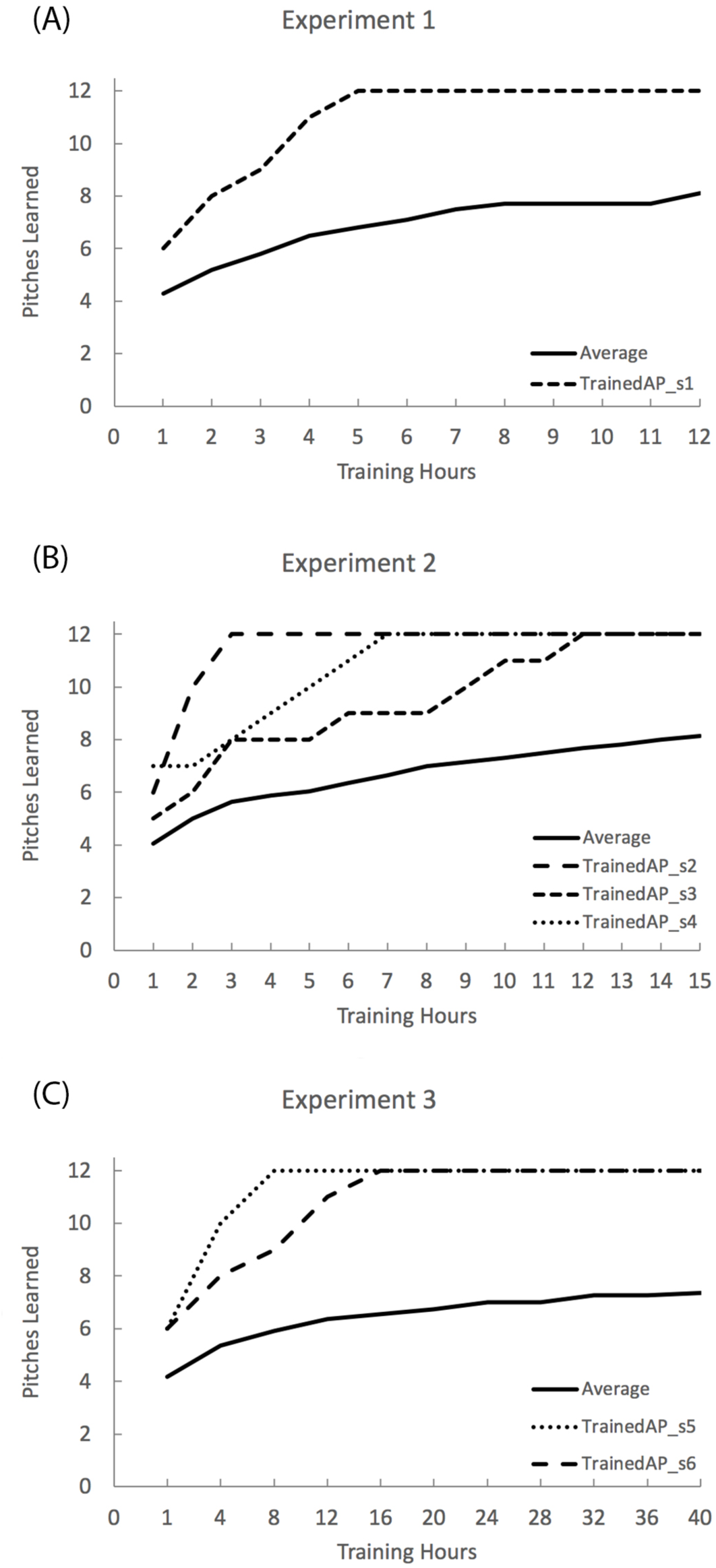
The number of pitches learned in the course of training in the three experiments. Number of pitches learned was defined as the number of pitches included at the highest passed levels at 90% accuracy with no feedback or scoring assistance in each individual. The solid line shows the average number of learned pitches across all individuals in each experiment. The other lines showed the course of training of the individuals who passed all levels of training in each experiment. TrainedAP_s6 in Experiment 3 learned to name 12 pitches at 90% accuracy with a subset of training tones (in one octave and one timbre only at level 74) at the end of the 16^th^ hour of training, but she actually passed all levels of training (level 80) during the 18^th^ hour of training.

### Generalization & sustainability of AP learning

The pitch-naming performance improved and was similar between trained and untrained tones (Figure 3A). A 2 x 3 ANOVA with Prepost (pretest / posttest) and Stimulus Type (octave & timbre trained / octave trained & timbre untrained / octave untrained & timbre trained) as factors on pitch naming error revealed a significant main effect of Prepost, *F*(1,9) = 30.3, *p* < .001, *η*_p_^2^ = .771, with a smaller pitch naming error at posttest than pretest. No other main effect or interaction was observed, *p*s > .15, i.e., we did not observe any difference between the naming performance of tones in trained or untrained timbres and octaves (*η*_p_^2^ of the main effect of Prepost was .846, .638, .485 for octave & timbre trained, octave trained & timbre untrained, octave untrained & timbre trained conditions respectively).

**Figure 3.**
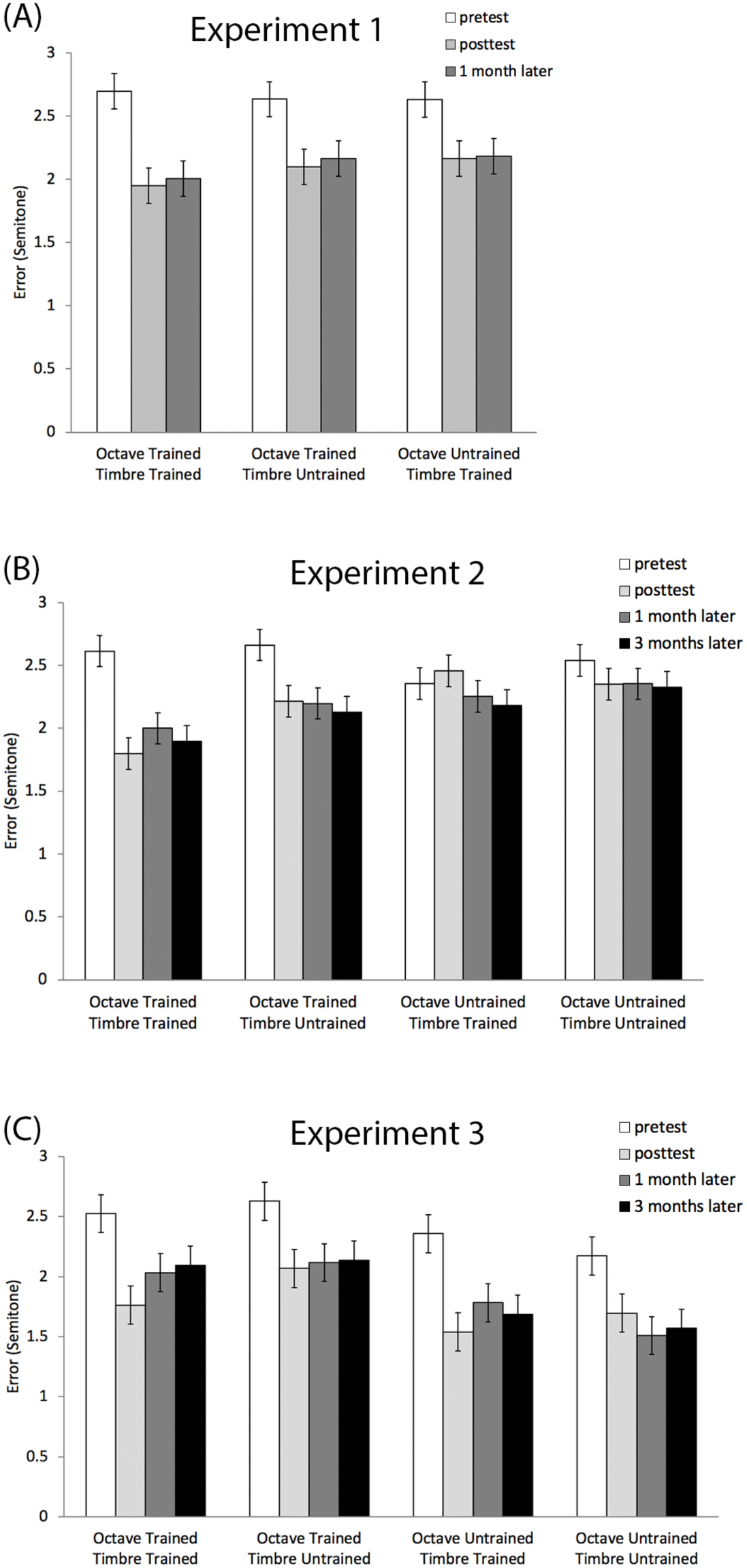
Generalization of learning to different octaves and timbre in the three experiments. Pitch naming error was defined as the deviation in semitone from the correct response. Error bars represent 95% CIs for the interaction effect between Prepost and Stimulus Type in Experiment 1 (A), for the interaction effect between Prepost, Octave and Timbre in Experiments 2 and 3 (B-C).

To check if the improvement sustained for a month, a one-way ANOVA was performed with Prepost (pretest / posttest / a month later) on pitch naming error with the trained tones^2^. It revealed a significant main effect of Prepost, *F*(2,16) = 18.8, *p* < .001, *η*_p_^2^ = .701. Post-hoc Scheffé tests (*p*<.05) showed that the pitch naming error reduced after training and remained similar a month later, *p* > .73 (*η*_p_^2^ of the main effect of Prepost between the pretest and the immediate posttest was .840 and that between the pretest and the posttest a month later was .675).

## Discussion

After the 12-hour AP training, all participants improved their pitch naming performance. On average, they were able to name 8.1 pitches accurately. In particular, one of the participants was able to name all of the twelve pitches at 90% accuracy without externally provided reference tones. This level of AP performance was representative of and comparable to that of ‘AP possessors’ as defined in the literature. This indicates that AP acquisition is possible in adulthood, and a 12-hour training protocol was sufficient for at least one of the participants to acquire AP.

In addition, the characteristics of AP learning matched well with that of perceptual learning (Fahle & Poggio, 2002; Goldstone, 1998). Specifically, AP performance improved after training, and the improved performance did not differ between tones in trained or untrained timbres and octaves, suggesting that the AP learning generalized to untrained octaves and timbres. Also, the AP performance was similar right after training and a month later, suggesting that the AP learning sustained for a month. Overall, the AP learning corresponded well with classic characteristics of perceptual learning in terms of performance enhancement, generalization and sustainability.

While we did not include any control group to control for the potential learning effect during repeated testing, the observed improvement during the test for generalization was unlikely explained by repeated testing per se. In particular, AP is notoriously famous for its difficulty to acquire in the literature, and the average progress per hour of training (where at least hundreds of trials were involved) was relatively slow (Figure 2A). Therefore simply performing the test for generalization twice can unlikely cause the substantial and sustained AP improvement after training.

## Experiment 2

In Experiment 2, we aimed to replicate the feasibility of acquiring AP in adulthood and further characterize such AP learning. First, we tested the robustness of AP acquisition in adulthood by using a different set of training protocol, including a different set of training tones, training tasks, training duration and design (Table 1 and Figure 1). Second, we asked whether training with a smaller set of stimuli, i.e., tones in one octave and one timbre only, would lead to higher specificity in AP learning since it covers the dimensions of octave and timbre more specifically in the psychological space (Nosofsky, 1986, 1987; Palmeri & Gauthier, 2004; Y. K. Wong et al., 2011). Third, we also examined whether musicians benefit from the training more than non-musicians due to their prior musical training.

## Methods

### Participants

Twenty-two participants, including ten musicians and twelve non-musicians, were recruited at City University of Hong Kong and completed the training. The musicians included 4 males and 6 females and were 23.5 years old on average (*SD* = 3.34). They were trained in music for 6-14 years, with the major instrument being piano (*N* = 9) and guitar (*N* = 1). The non-musicians included 5 males and 7 females and were 21.8 years old on average (*SD* = 1.27). Two additional musicians dropped out of the training soon after participating because they could not commit to the whole training. The sample size per group was determined by matching that of Experiment 1. All participants filled out a similar questionnaire on musical training background similar to that of Experiment 1. Participants received monetary compensation for the training and testing, with additional bonuses for passing each level of training. Informed consent was obtained according to the Ethics Committee of City University of Hong Kong.

### Materials

In Experiment 2, 72 tones in timbres of synthetic, piano and violin tones from octaves four to five were used. The tones were identical to that used in Experiment 1. An additional glissando clip, which was often perceived as an endless glissando travelling continuously from a high pitch to a low pitch (Deutsch, 1995; Deutsch, Hamaoui, & Henthorn, 2007), was used to further interfere any existing auditory memory of tones before AP testing (see below).

### Absolute pitch training

The training included 12 synthetic tones from octave 4. Similar to Experiment 1, the training was gamified and structured with different levels. If participants achieved 90% accuracy for a certain level, they would proceed to the next level; otherwise they would stay at the same level. The training was completed by finishing 15 hours of training or by passing all 30 levels with 90% accuracy. Participants finished one hour of training per day. They were trained four days per week and finished the training in four weeks.

The 30-level training protocol was organized into ten 3-level parts with an increasing number of pitches (from 3 pitches in the first 3 levels to 12 pitches in the last 3 levels; Figure 1C). Within each 3-level part, the first level involved a verification task, in which participants judged by key press whether the pitch of the presented tone matched with a pitch name shown on the screen (‘y’ for match and ‘n’ for mismatch). Trial-by-trial feedback was provided. The second level was a naming task, in which participants named the pitch of the tones by key press based on an image showing the pitch-to-key mapping (Figure 1A). Trial-by-trial feedback was provided. The third level was a similar naming task without trial-by-trial feedback. If participants achieved 90% accuracy with a specific set of pitches at the third level, a new pitch was introduced into the training set, and they went through the three-level part again. While participants generally stayed at the same level if they did not pass it, for this third level, upon failure in three consecutive attempts, they would be moved back to the second level. This allowed participants to re-learn the materials with the assistance of trial-by-trial feedback in case they were not ready for no-feedback training. The number of trials increased from 12 trials per level to 30 trials per level gradually, with two trials added every time a new pitch was introduced. This allowed the blocks of trials to better represent and capture the increasing number of pitches included in the training set. At each level, the tones were distributed as evenly as possible among the training pitches, octaves and timbres. Semitone errors were considered errors in the training. Before each level, participants were allowed to freely listen to sample tones of the training pitches as many times as they preferred before proceeding to the training. Each training session lasted for an hour, in which individual participants might have finished different numbers of training trials depending on their pace of learning (e.g., the amount of time spent on the training trials or on sample tone listening). A similar system of three-point special trials and token accumulation applied as in Experiment 1.

Similar to Experiment 1, the no-feedback levels were designed to serve as mini milestones for participants’ AP performance at 90% accuracy. To further destroy any existing auditory memory trace of tones before testing, a 15s glissando clip was played after listening to the sample tones and before the start of the no-feedback levels. This should have effectively interfered with any auditory memory trace of tones since a 15-s latency with ineffective rehearsal (because of interfering tasks or stimuli) has been shown to result in very low accuracy during recall (10% or lower; Peterson & Peterson, 1959; see also the discussion in Takeuchi & Hulse, 1993; and Wengenroth et al., 2013). This further minimized the possibility that participants performed the no-feedback levels with the assistance of external reference tones.

### Test for generalization

Similar to Experiment 1, the test for generalization was performed before, within three days after training, a month later and three months later to examine how well the pitch-naming abilities generalized to untrained octaves and timbres. Seventy-two tones were used. The tones were either in octaves 4 to 5, in which octave 4 was trained and octave 5 was untrained. Also, the tones were either in three timbres, with synthetic tones as a trained timbre, and piano and violin tones as untrained timbres. The test was administered with four conditions, either with trained octave and timbre, trained octave and untrained timbre, untrained octave and trained timbre, or untrained octave and timbre. A similar pitch-naming task was used as in Experiment 1. There were 144 trials presented in randomized order, with each tone presented twice. Ten practice trials were provided with feedback before testing. The 15s glissando tone was presented after the practice with feedback and before the testing to minimize any auditory working memory of tones that were previously heard (Peterson & Peterson, 1959; Takeuchi & Hulse, 1993; Wengenroth et al., 2013). Pitch-naming error, i.e., the average error in semitone of participants’ response in comparison with the correct response, was used as the dependent measure.

## Results

### Learning of AP

In general, participants made substantial progress in learning to name pitches. At the end of training, the musicians and non-musicians were able to name on average 9.4 (range = 7-12; SD = 2.12) and 7.2 pitches (range = 5-9; SD = 1.34) at 90% accuracy respectively (Figure 2B). A one-way ANOVA with Group (musician / non-musician) on number of learned pitches by the end of training revealed a significant main effect of Group, *F*(1,20) = 9.06, *p* = .007, *η*_p_^2^ = .312. This shows that musicians had a larger progress in learning AP than non-musicians under the same training regime.

Importantly, three out of the twenty-two participants passed all levels of training, meaning that they were able to name all of the twelve pitches at 90% accuracy without externally provided reference tones, suggesting that they have acquired AP through perceptual learning in adulthood, with their AP performance comparable to that of real-world ‘AP possessors’ (for definition see Methods in Experiment 1).

### Generalization & sustainability of AP learning

Pitch-naming performance improved and the improvement was larger for the trained than untrained tones (Figure 3B). A 2 x 2 x 2 x 2 ANOVA with Group (musician / non-musician), Prepost (pretest / posttest), Octave (trained / untrained) and Timbre (trained / untrained) as factors on pitch naming error revealed a significant main effect of Prepost, *F*(1,20) = 25.6, *p* < .001, *η*_p_^2^ = .561, as the pitch naming error reduced after training. There were significant interactions between Prepost and Octave, *F*(1,20) = 11.4, *p* = .003, *η*_p_^2^ = .363, and between Prepost, Octave and Timbre, *F*(1,20) = 13.8, *p* = .001, *η*_p_^2^ = .408. Post-hoc Scheffé test (*p* < .05) showed that the precision of pitch naming was higher for tones with the trained octave and timbre than other tones only after training. The interaction between Prepost and Group was significant, *F*(1,20) = 4.87, *p* = .039, *η*_p_^2^ = .196, in which both groups showed better performance at posttest than pretest, with a larger improvement in musicians (*M*_*pretest*_ = 2.39, *SD* = .64; *M*_*posttest*_ = 1.88, *SD* = .76; *η*_p_^2^ for the main effect of Prepost = .618) than non-musicians (*M*_*pretest*_ = 2.67, *SD* = .36; *M*_*posttest*_ = 2.47, *SD* = .53; *η*_p_^2^ for the main effect of Prepost = .470). The main effect of Timbre was significant, *F*(1,20) = 8.93, *p* = .007, *η*_p_^2^ = .309, which interacted with Group, *F*(1,20) = 5.29, *p* = .032, *η*_p_^2^ = .209. Post-hoc Scheffé test (*p* < .05) showed that non-musicians performed better with the trained timbre than the untrained timbre, while musicians performed similarly with the two timbres. The four-way interaction did not reach significance (*F* < 1).

To check if the improvement sustained, a two-way ANOVA with Group (musician / non-musician) x Prepost (pretest / posttest / 1 month later / 3 months later) was performed on pitch naming error with the trained tones only^3^. It revealed a significant main effect of Prepost, *F*(3,54) = 17.4, *p* < .001, *η*_p_^2^ = .492. Post-hoc Scheffé tests (*p* < .05) showed that pitch naming error was smaller at posttest than pretest, and stayed similar one and three months later (*η*_p_^2^ of the main effect of Prepost between the pretest and each of the subsequent tests was .614, .636 and .554 for the posttest, 1 month later and 3 months later respectively), suggesting that the AP improvement sustained for at least three months. There was a trend that experts performed better than novices in general (*p* = .09), but this effect did not interact with Prepost (*F* < 1).

## Discussion

In Experiment 2, we replicated the feasibility of acquiring AP in adulthood and further characterized AP learning. First, three adults successfully acquired AP using a different set of training protocols, suggesting that AP acquisition in adulthood is not a result of a certain specific type of training protocol used in Experiment 1, but can be generally observed with different perceptual learning paradigms. Second, we replicated the perceptual learning nature of AP learning in adulthood in terms of performance enhancement and sustainable improvement for at least three months. In terms of generalization of learning, the AP learning in this experiment was less generalizable to tones in untrained timbres and octaves than in Experiment 1, because here the AP training only covered training tones from a single octave and timbre. This is consistent with the hypothesis that the degree of learning specificity and generalization depends on the psychological multidimensional space of the training and testing stimuli (Nosofsky, 1986, 1987; Palmeri & Gauthier, 2004; Y. K. Wong et al., 2011). The higher improvement for the trained tones was also inconsistent with the possibility that the observed improvement during the tests for generalization was caused by repeated testing per se, since any improvement caused by repeated testing should be comparable in all conditions that were similarly exposed. Finally, musicians learned to name more pitches than non-musicians during the training, and showed a larger improvement than novices in the precision of pitch naming during the test for generalization, suggesting that previous exposure to musical tones may facilitate AP acquisition. One caveat is that we recruited real-world musicians and non-musicians. Without manipulating musical experience, we could not exclude the possibility that pre-existing differences in the two groups that were irrelevant to prior music exposure (e.g., higher motivation in musicians) might explain the group difference in AP learning.

## Experiment 3

In Experiment 3, we further asked whether AP training in adulthood is feasible outside of the laboratory. Participants performed an online AP training anywhere with a stable Internet connection. They each finished 40 hours of training in their own pace within an eight-week period.

## Methods

### Participants

Eleven participants were recruited at City University of Hong Kong and completed the training. They included 4 males and 7 females and were 20.1 years old on average (*SD* = 0.83). Six of them have received musical training for 3 to 15 years, with the major instrument being piano (N = 3), violin (N = 2) and flute (N = 1). Five were non-musicians who were not formally trained with music or had brief training that lasted for less than a year. They performed the Distorted Tunes Test for tone-deaf screening at pretest (available at https://www.nidcd.nih.gov/tunestest/test-your-sense-pitch) so as to ensure that none of the participants would encounter difficulty in acquiring AP simply because of deficits in perceiving pitch in general. All participants performed the task satisfactorily with no suspected deficit with detecting distorted tunes (accuracy ranged from 73.1% to 100%, *M* = 88.2%, *SD* = 0.088). One additional participant was excluded from the training with highly accurate pitch naming precision during the pretest (with an average of 0.72 semitones from correct responses), leaving the training unnecessary. One additional musician was excluded from data analyses because she showed highly inconsistent and uninterpretable training performance^4^. The sample size was determined to match that of Experiments 1 and 2. All participants filled out a similar questionnaire on musical training background similar to that of Experiment 1. Participants received monetary compensation for the training and testing, with additional bonuses for passing each level of training. Informed consent was obtained according to the Ethics Committee of City University of Hong Kong.

### Materials

In Experiment 3, 120 tones were used, including that in timbres of synthetic and piano tones in octaves four to six, and that in timbres of violin and clarinet in octaves four to five. The synthetic, piano and violin tones were identical to that used in Experiments 1 and 2. The clarinet tones were downloaded from online sound library (http://newt.phys.unsw.edu.au/music/clarinet/index.html) and edited in an identical way as the other tones. The same glissando clip in Experiment 2 was also used in this experiment.

### Absolute pitch training

This training was administered online such that participants could be trained anywhere with stable Internet connection. It included piano, violin and clarinet tones in octaves 4 and 5. A pitch-naming task was used. During each trial, an isolated tone was presented for 1s. Then, the pitch-to-key mapping was shown (Figure 1A). Participants were required to name the pitch of the presented tone by mouse clicking within 5s.

Similar to Experiments 1 and 2, the training was gamified and structured with different levels. If participants achieved 90% accuracy for a certain level, they would proceed to the next level; otherwise they would stay at the same level. The training was completed by finishing 40 hours of training or by passing all 80 levels with 90% accuracy. The training time of the participants was counted into the total training time in units of 15 minutes (i.e., 17 minutes training was taken as 15 minutes, and 14 minutes training as 0 minute). To minimize any idling time, the session for listening to sample tones before each level would be ended with 10s of inactivity. The program would automatically log out if there was inactivity for 3 minutes. They were trained for an average of five hours per week and finished the training in about eight weeks.

Similar to Experiment 1, the training protocol involved 80 levels and was organized into ten 8-level parts with an increasing number of pitches (from 3 pitches in the first 8 levels to 12 pitches in the last 8 levels; Figure 1D). Each eight-level part consisted of four types of levels, which included tones that were progressively richer in timbres and octaves. Each of the four types of levels was repeated twice, once with feedback provided, and then once without feedback. For example, participants began the training with three pitches (E, F and F#). At levels 1-2, piano tones in these three pitches in octave 4 were included, with feedback provided at level 1 and then without feedback at level 2. At levels 3-4, piano tones in both octave 4 and 5 were included with feedback and then without feedback. At levels 5-6, piano, violin and clarinet tones in octave 4 were included with feedback and then without feedback. At levels 7-8, piano, violin and clarinet tones in octave 4 and 5 were included with feedback and then without feedback. At the no-feedback levels, participants were not provided with any external feedback of the correctness of the tones. If they achieved 90% accuracy at the 8^th^ level, a new pitch was added into the training set, with which they went through the same eight-level part again. The number of trials increased from 12 trials per level to 30 trials per level gradually, with two trials added every time a new pitch was introduced. This allowed the training to better represent and capture the increasing number of pitches included in the training set. At each level, the tones were distributed as evenly as possible among the training pitches, octaves and timbres. Semitone errors were considered errors in the training. Similar to Experiment 2, participants were allowed to freely listen to sample tones of the training pitches as many times as they preferred before proceeding to each level of the training. The same glissando clip was played after listening to the sample tones and before the start of the no-feedback levels so as to destroy any existing auditory memory trace of previously heard tones. Also, the same system of tokens and three-point special trials were adopted in this study. Importantly, the special trials did not appear and could not be initiated during the no-feedback levels, similar to Experiments 1 and 2. This ensured that participants performed the no-feedback levels without any scoring assistance.

To promote better learning, a summary table of the participant’s performance was provided at the end of each level. The table showed the accuracy for each pitch and listed out the wrong answers provided for each pitch (if any). This information enabled participants to better evaluate the type of errors they made. Also, when the training set included 5 pitches or more, an additional compulsory exercise was introduced after every 10 attempts of passing the levels. The exercise would include 20 trials that centered around the worst-performed pitch in the past 10 attempted levels. The timbre and octave of the tones would follow the level right before this exercise. This was designed to further help participants focus on the pitches that needed most training.

### Test for generalization

The test for generalization was performed in the laboratory before and within three days after training to examine how well the pitch-naming abilities acquired online generalized to trained and untrained octaves and timbres in the laboratory setting. In this test, 72 tones in octaves 4 to 6 (C4 to B6) were used, in which octaves 4 and 5 were trained, and octave 6 untrained. Two timbres were included, with piano tones as the trained timbre, and synthetic tones as the untrained timbre. A similar pitch naming task as in Experiments 1 and 2 was used. During each trial, a tone was presented for 1s. Then an image showing the pitch-to-key mapping appeared. Participants were required to name the pitch of the presented tone by key press within 5s. The trials were blocked in four conditions, either with a trained or untrained octave crossed with a trained or untrained timbre. There were 192 trials in total, with 48 trials in each condition. The trials within each block were randomized. Similar to the test in Experiment 2, ten practice trials were provided with feedback before testing. A 15s glissando tone was presented after the practice but before the testing to destroy any existing auditory working memory of tones that were previously heard (Peterson & Peterson, 1959; Takeuchi & Hulse, 1993; Wengenroth et al., 2013). Pitch-naming error, i.e., the average error in semitone of participants’ response in comparison with the correct response, was used as the dependent measure.

## Results

### Learning of AP

In general, the participants made substantial progress in learning to name pitches. At the end of training, they were able to name on average 7.4 pitches (range = 5-12; SD = 2.46) at 90% accuracy (Figure 2C). Importantly, two out of the eleven participants passed all levels of training, meaning that they were able to name all of the twelve pitches at 90% accuracy without any externally provided reference tones. These suggest that they have acquired AP through perceptual learning in adulthood, with a performance level comparable to that of real-world ‘AP possessors’ (for definition see Methods in Experiment 1).

### Generalization & sustainability of AP learning

The online AP training improved pitch-naming performance in the laboratory, and the improvement generalized well to untrained octaves and less so to untrained timbres (Figure 3C). A 2 x 2 x 2 ANOVA with Prepost (pretest / posttest), Octave (trained / untrained) and Timbre (trained / untrained) as factors on pitch naming error revealed a significant main effect of Prepost, *F*(1,10) = 15.0, *p* = .003, *η*_p_^2^ = .599, as pitch naming error reduced after training. The interaction between Prepost and Timbre was significant, *F*(1,10) = 7.15, *p* = .023, *η*_p_^2^ = .417. Post-hoc Scheffé tests (*p* < .05) showed that the pitch naming error was similar for trained and untrained timbres before training. While the error of both types of timbres reduced at posttest, the error was smaller for trained timbres than untrained timbres after training. The main effect of Octave was significant, *F*(1,10) = 15.8, *p* = .003, *η*_p_^2^ = .609, as the pitch naming error was smaller for the untrained than trained octave in general.

However, the interaction between Prepost and Octave did not reach significance (*F* < 1), meaning that we did not observe any difference in the degree of AP learning for trained and untrained octaves (*η*_p_^2^ for the main effect of Prepost was .643 and .500 for the trained and untrained octaves respectively). The three-way interaction did not reach significance (*F* < 1).

For the sustainability of the improvement, a one-way ANOVA with Prepost (pretest / posttest / 1 month later / 3 months later) on pitch naming error with the trained tones revealed a main effect of Prepost, *F*(3,27) = 10.6, *p* < .001, *η*_p_^2^ = .541^5^. Post-hoc Scheffé tests (*p* < .05) showed that pitch naming error reduced after training, and stayed similar at all subsequent posttests (*η*_p_^2^ of the main effect of Prepost between the pretest and each of the subsequent tests was .676, .518 and .522 for the posttest, 1 month later and 3 months later respectively), suggesting that the improvement sustained for at least three months.

## Discussion

In Experiment 3, we further replicated the finding that AP can be acquired in adulthood. Using a 40-hour training on the Internet at anywhere at one’s own pace, two adults were able to name tones in all twelve pitches at 90% accuracy without external reference tones, and thus have successfully acquired AP through perceptual learning in adulthood. This demonstrates that AP training is feasible outside of the laboratory.

The characteristics of AP learning matched with that of perceptual learning well. Using tones in multiple timbres and octaves, naming accuracy improved for both trained and untrained tones, suggesting that AP learning generalized to untrained tones similar to findings of Experiment 1. The improvement for trained timbres was larger than that of untrained timbres, while the improvement for trained and untrained octaves was similar. This indicated that the generalization of learning was more complete for octave than timbre. The AP improvement sustained for at least three months. Overall, the performance enhancement, generalization and sustainability correspond well with that of the perceptual learning literature (Fahle & Poggio, 2002; Goldstone, 1998; Y. K. Wong et al., 2011).

## Overall Analyses of Experiments 1-3

### Characterizing individual progress of AP learning

Is it the case that one can either acquire AP, or be completely unable to improve this ability? We did observe such a continuous distribution of learning progress in this study (Figure 4), without a considerable number of participants acquiring every step between 5 to 12 pitches. If our training could further extend in time, it is well possible that more participants might be able to learn all of the twelve pitches (e.g., one participant who learned 11 pitches and two participants who learned 10 pitches by the end of the training). In general, our results were more consistent with the continuous view of AP ability (Bermudez et al., 2009; Levitin & Rogers, 2005) than the dichotomous view (Athos et al., 2007; W. D. Ward, 1999; Zatorre, 2003).

**Figure 4.**
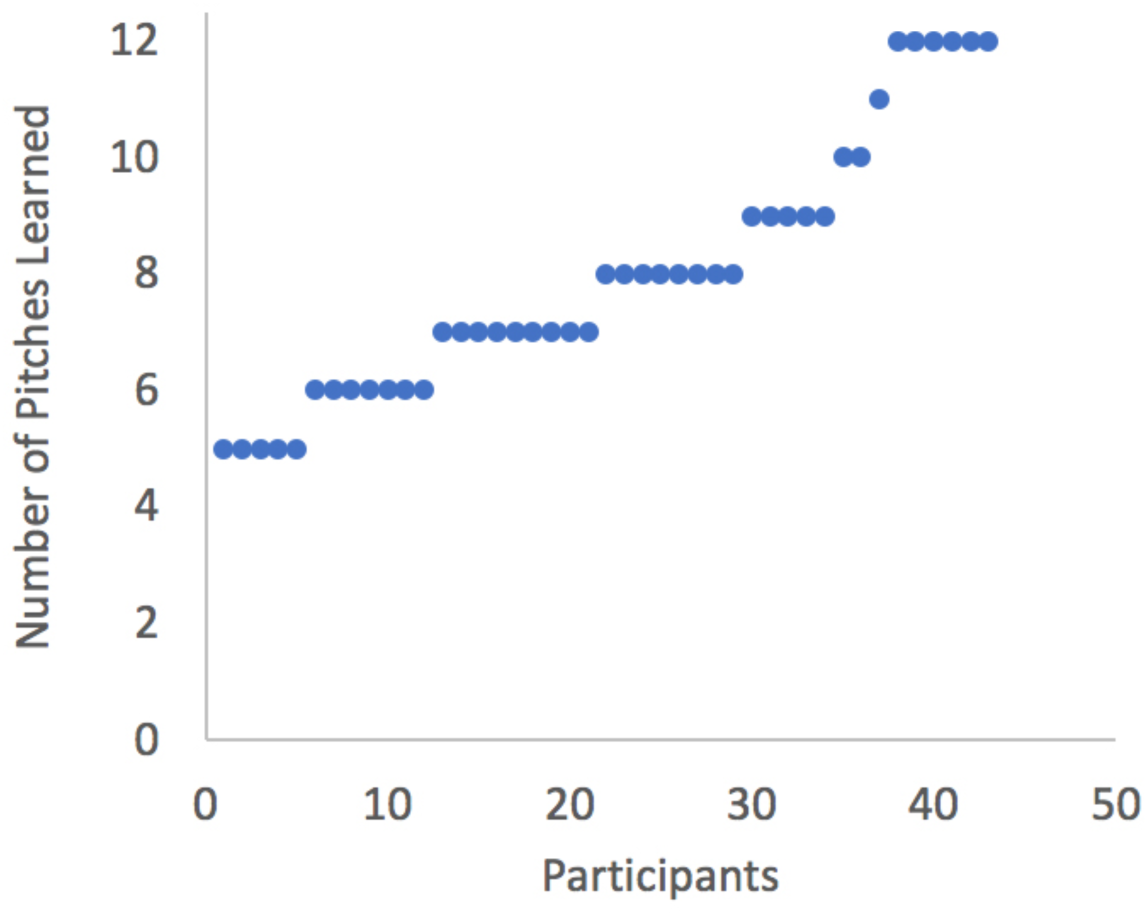
The number of pitches learned by the end of training, with the participants ranked from low to high number of learned pitches.

### Characterizing the trained AP

Despite the continuous training progress across participants, it is still informative to further examine the participants who learned to name all 12 pitches. Across the three experiments, we identified six out of 43 (14.0%) participants who finished all the training levels and thus learned to name all 12 pitches. An obvious question concerns whether these successfully trained participants were simply ‘AP possessors’ before training. Two aspects of our data indicate that they were not. One is that they spent much longer time to pass all of the training levels than one would expect if they were already ‘AP possessors’ before training. Specifically, the AP training should be highly straightforward for ‘AP possessors’, and thus passing each of the levels should be easy with one or two attempts. If so, they should be able to pass the training well within the first hour of training. However, they required 7.83 hours on average to pass the training (SD = 5.23; range: 3 to 18; Figure 2), which was unreasonably long if they were ‘AP possessors’. Second, their pretest performance of AP naming for the training tones was low (*M*_*error*_ = 1.97 semitones; *SD*_*error*_ = 0.47). This performance level, even considering the fact that the timbre and octave used may not be the most familiar to these individuals, would still be rarely considered typical of ‘AP possessors’ among the published studies that defined ‘AP possessors’ based on AP performance measures. Therefore, the pretest performance and training progress of these successfully trained participants did not support the idea that they were ‘AP possessors’ before training.

Next, was the music training background of the successfully trained participants different from the other musicians? In terms of musical achievement (indicated by the highest passed level of ABRSM^6^) and the age of onset of musical training, the successfully trained participants were well within 3 SD of the mean of the other musicians participated in this study (range = -.936 to 1.63 SD and -.374 to −1.10 SD from the mean of other musicians for ABRSM and for age of onset respectively). Therefore we did not see clear evidence that the successfully trained participants had a different music training background from other musicians.

In the literature, it is well known that ‘AP possessors’ suffer from performance decrement with unfamiliar timbres and octaves (Levitin & Rogers, 2005; Takeuchi & Hulse, 1993; Y. K. Wong & Wong, 2014). In our test for generalization, we included a large proportion of untrained tones with varied timbres and octaves, ranging from 66.7% to 75% of the trials, and the untrained tones were often intermixed with the trained tones. Was the trained AP ability also affected by the exposure to a large proportion of untrained tones? To address this question, we examined the AP performance of the trained tones during the posttest right after training for these six individuals. A one-way ANOVA with Prepost (pretest / posttest) on pitch naming error with the trained tones revealed a main effect of Prepost, *F*(1,5) = 16.4, *p* = .010, *η*_p_^2^ = .766, indicating that the pitch naming error was smaller at posttest (*M*_*error*_ = .925 semitones; *SD*_*error*_ = 0.45) than pretest (*M*_*error*_ = 1.97 semitones; *SD*_*error*_ = 0.47). There was however a wide range of pitch naming errors across individuals (range: .375 – 1.58 semitones), suggesting that the trained AP of some individuals were highly susceptible to the exposure to a large proportion of unfamiliar tones.

Finally, did the trained AP sustain for at least a month? A one-way ANOVA with Prepost (pretest / posttest / 1 month later)^7^ on pitch naming error with the trained tones revealed a main effect of Prepost, *F*(2,10) = 17.1, *p* < .001, *η*_p_^2^ = .773. Post-hoc LSD test (*p* < .05) showed that the pitch naming error was smaller at posttest than pretest and remained similar a month later (*M*_*error*_ = .958 semitones; *SD*_*error*_ = 0.50; *η*_p_^2^ of the main effect of Prepost between the pretest and each of the subsequent tests was .766 and .848 for the posttest and 1 month later respectively), suggesting that the trained AP sustained for at least a month.

### Comparing different training paradigms

The three training paradigms used in the three experiments differed in many ways, including the training stimuli, duration, the venue of training, and the design of the training progression. While it is difficult to pinpoint the contributions of specific training parameters, we can at least view these training paradigms as different packages and ask one question: Was any of the training paradigms more effective than the others in general? A one-way ANOVA with Experiment (1 /2 /3) as a between-subject factor was performed on the number of learned pitches at the end of the training, with the number of learned pitches defined by the number of pitches included at the highest passed level without feedback. The main effect of Experiment did not reach significance (*F* < 1; *η*_p_^2^ of the main effect of Prepost was .852, .680, .593 for Experiment 1, 2 and 3 respectively), suggesting that the AP learning was not significantly different between training paradigms.

Apart from the progress during training, we can also compare the degree of generalization in the three training paradigms. The results from the three experiments suggested that the degree of generalization varied between the experiments, with more generalization of learning in Experiment 1 and 3 (in which more timbre and octave were involved during training) than in Experiment 2. A 3×2×3 ANOVA with Experiment (1 /2 /3) as a between-subject factor, and Prepost (pretest / posttest) and Stimulus Type (octave & timbre trained / octave trained & timbre untrained / octave untrained & timbre trained) as two within-subject factors was performed on pitch naming error. Critically, the three-way interaction between Experiment, Prepost and Stimulus Type was significant, *F*(4,82) = 3.27, *p* = .015, *η*_p_^2^ = .138. Post-hoc Scheffé test (*p* < .05) showed that performance in all stimulus conditions were similar within each experiment at pretest. At posttest, however, pitch naming error was similar between the trained and untrained tones for Experiment 1 and 3, while the pitch naming error was smaller for the trained than untrained tones in Experiment 2. This confirmed that the generalization of learning to untrained tones was higher in Experiment 1 and 3 than in Experiment 2.

## General Discussion

The observations that AP rarely occurs in the population and is difficult to learn in adulthood have led to the belief that AP is only possible for few selected individuals - those with particular genetic makeup and musical training within a critical period in early childhood. Our study provides the first empirical evidence that pitch naming can improve considerably in adults in both laboratory and online settings. There was a continuum of training progress among the participants in terms of the number of pitches learned. Importantly, 14% of the participants (6 out of 43) successfully acquired AP within the training period, with performance levels comparable to that of ‘AP possessors as defined in the literature. This proportion of training success is unprecedented in the literature of AP, suggesting that AP continues to be learnable in adulthood.

The pattern of AP learning is consistent with that of the perceptual learning literature. AP performance improved for all participants after training. In particular, the learning effects sustained for at least one to three months and generalized to untrained tones to an extent depending on the specificity of the training set. When the training set included tones from more octaves and timbres (e.g., Experiments 1 & 3), AP learning generalized better to untrained tones, whereas the learning was more specific to the trained tones when the training targeted on a smaller set of tones (e.g., Experiment 2). The specificity of learning as a function of the psychological multidimensional space during training and testing is consistent with findings in perceptual learning studies (Fahle & Poggio, 2002; Goldstone, 1998; Y. K. Wong et al., 2011). We did not observe any difference in the number of pitches learned in the three experiments that involved different training protocols, tasks, stimuli and durations. This suggests that AP learning in adulthood can be achieved with a wide range of training regimes, similar to perceptual learning in general (Fahle & Poggio, 2002).

### Did the training lead to ‘genuine AP’?

In this study, AP was empirically defined based on performance in a pitch-naming task. It is almost the only empirical measure used in the literature to distinguish between ‘AP possessors’ and others, based on which insights on absolute pitch processing have been obtained. Also, the criterion used for AP acquisition (90% accuracy in naming all 12 pitches, with one semitone error regarded as incorrect) is more stringent than that used in the majority of empirical studies thus far. Based on the review of the literature (see Methods of Experiment 1), participants passing this criterion would also be regarded ‘AP possessors’ in over 80% of the studies. We therefore deem it appropriate to use this operational definition to assess AP acquisition.

One may question whether the participants may have used relative pitch strategies to facilitate their AP performance during training and testing, given that sample tones and feedback were provided for certain training trials, and practice trials during pre- and posttests. It should be noted that a 15s glissando clip was introduced after listening to the sample tones in training, before the start of the no-feedback levels in training, and after the feedback-provided practice trials in pre- and posttests. This has been shown to be effective in interfering with and minimizing the auditory memory trace left by any sample tones heard (Peterson & Peterson, 1959; Wengenroth et al., 2013). If, even with the glissando clip, the sample tones still served effectively as externally provided reference, then the AP training and testing should essentially become relative pitch training and testing. This should have resulted in much better performance in our training and testing, which was not the case – the pretest performance was chance-like (Figure 3) and the training progressed very slowly on average (Figure 2).

Still, an alternative interpretation of the current finding is that the participants simply improved their accuracy in naming pitches through training, but were yet to acquire ‘genuine’ AP. With the relatively long time windows allowed for response (5s for choosing one of twelve options by key pressing or mouse clicking), participant could have generated reference tones internally in a trial and use it for subsequent trials. In other words, participants could have learned to represent some of the pitches in an absolute manner while naming other pitches by comparing them to the internally generated references. While one can make such relative pitch comparisons more difficult by introducing a large pitch interval between adjacent trials, it was impossible in the current study given the pitches presented were generally selected from narrow ranges (within one to two octaves). If comparison with internal references is indeed what participants in general learned to do, then the ability acquired would be considered by some researchers as ‘quasi-AP’ (despite their accurate AP performance without relying on external reference tones), rather than the ‘genuine AP’ involving the representation of every pitch in an absolute manner (e.g., Levitin & Rogers, 2005).

Regarding this alternative interpretation, two points are worth noting. First, the majority of empirical research has identified individuals as ‘AP possessors’ based on performance alone, without regarding to the underlying representation of pitch through which the high performance level is achieved. Our findings here would therefore be relevant and informative to the research field in general. Second, it is certainly interesting and important to ask whether those identified as ‘AP possessors’ in most empirical studies indeed represent every pitch absolutely. However, to date there is not an agreed upon way to ascertain whether an individual perceives every pitch in an absolute manner or not. One could perhaps use fast responses to infer automatic pitch naming in an absolute manner. But the search for the threshold response time could be elusive - given that many factors would greatly affect response times, such as the stimulus set, the types of response (voice, key press, mouse click, etc.) and age, how fast is ‘sufficiently fast’ to represent ‘automatic’ pitch naming? Without an objective way to reveal the underlying representation of pitch of ‘AP possessors’, the assumption of ‘genuine AP’, as defined by representing every pitch absolutely, remains an untestable hypothesis. Still, future research with a larger sample of individuals with and without AP in the same study can examine if there are any qualitative and quantitative differences in terms of response time, accuracy, brain areas involved, etc. It will be then possible to investigate if the trained AP possessors recruit similar cognitive and neural mechanisms as the very best natural ‘AP possessors’.

### Challenges to explaining AP occurrence with genes and the critical period

Our finding of adults acquiring AP across all three experiments challenges the strong form of the genetic accounts of AP development that only a very small percentage of the population possesses the rare gene(s) that enables AP acquisition. Within our training duration, 14% of the participants acquired AP, which was likely a lower bound of the percentage of training success in adulthood. If our training could further extend in time, it is well possible that more participants might be able to acquire AP. Even for other participants, we observed a continuous range of AP abilities in terms of the number of learned pitches by the end of training and the pitch-naming error either before or after training. Together with the finding that AP exists in everybody to some extent before training (Halpern, 1989; Levitin, 1994; Schellenberg & Trehub, 2003), our results are more consistent with the continuous view (Bermudez et al., 2009; Levitin & Rogers, 2005) of the distribution of AP than the dichotomous view of AP abilities (Athos et al., 2007; W. D. Ward, 1999; Zatorre, 2003). Therefore, although we do not intend to deny the role of genes in AP acquisition, the current results suggest that any relevant genetic dispositions that *enable* AP acquisition should be a lot more prevalent in the population than previously assumed (Baharloo et al., 1998; Drayna, 2007).

Our finding also challenges the theoretical proposal that a critical period constrains the development of high-level cognitive abilities. This converges with findings in high-level vision (Daw, 1998) and musical development (Trainor, 2005) that high-level visual and musical abilities can continue to develop in adulthood. This is also consistent with the studies in language development (Flege et al., 1995; Singleton, 2001) and synesthesia (Bor et al., 2015), which show that acquisition of abilities comparable with native speakers and natural synesthetes beyond the critical period is possible. The critical period may instead mainly apply to the development of basic structures and functions, such as the anatomical development of the visual system based on the environmental inputs (Hubel & Wiesel, 1970; Simons & Land, 1987; Wiesel & Hubel, 1963; Zhang et al., 2002; but see Hooks & Chen 2007).

The participants in the current study were native speakers of Cantonese, which is a tonal language. In tonal languages, words that are different only in tones (pitch heights or contour) can have entirely different semantic meanings. For example, in Cantonese, the word ‘ma’ means ‘mother’ in tone one, ‘grandmother’ in tone four, ‘horse’ in tone five, and ‘to scold’ in tone six. It has been proposed that tonal language speakers learn to associate words with tonal templates that are absolute, precise and stable during the critical period of language development, which could later facilitate the development of absolute pitch ability (Deutsch et al., 2009; Deutsch, Henthorn, & Dolson, 2004; Deutsch, Henthorn, Marvin, & Xu, 2006). However, this hypothesis also assumes a speech-based critical period of AP acquisition for tonal language speakers:

“On this line of reasoning, it was hypothesized that in cultures where tone languages are spoken, infants generally acquire AP for the tones of their language during the critical period in which they acquire other features of their native language (Deutsch, 2002). When they reach the age at which they can begin musical training, they acquire AP for musical tones in the same way as they would acquire the tones of a second tone language (see, for example, Wayland and Guion, 2004). For such individuals, therefore, the acquisition of AP for musical tones should also be subject to a critical period.” (p.2399, Deutsch et al., 2009)

Therefore the hypothesis predicts that the acquisition of AP is impossible in adulthood even for tonal language speakers. Our findings challenge the existence of such a speech-based critical period of AP acquisition. Future studies may compare AP acquisition in tonal and non-tonal language speakers with various factors (musical background and experience, social-economic status, intelligence, etc.) controlled to understand the contribution of tonal language background on AP development.

### Understanding AP acquisition from the learning perspective

Integrating findings in the AP literature and the current study, we propose that the genesis of AP is better understood from a learning perspective. First, AP continues to be learnable from childhood (Crozier, 1997; Miyazaki & Ogawa, 2006; Sakakibara, 2014) to adulthood (the current study), from single tones (the current study; Brady, 1970; Meyer, 1899) to complex melodies and songs (Halpern, 1989; Levitin, 1994; Schellenberg & Trehub, 2003), and from musicians, non-musicians to the general public (Experiment 2 of the current study; Levitin, 1994; Schellenberg & Trehub, 2003). This suggests that AP is learnable in a wide range of ages, stimuli and individuals. Second, even for the ‘AP possessors’ that have stably acquired AP, they can still learn to fine-tune their AP representations based on recent environmental input (e.g., detuned music; Hedger et al., 2013), suggesting that their AP representations remains actively alterable instead of being ‘hard-wired’. Third, the learning perspective is consistent with the substantial evidence showing that AP is shaped by experience. For example, AP performance is better with the timbre of one’s own instrument (Takeuchi & Hulse, 1993), the highly exposed pitch like ‘A4’ (Levitin & Rogers, 2005; Takeuchi & Hulse, 1993), the multisensory testing context similar to one’s musical training (Y. K. Wong & Wong, 2014). Overall, AP is learnable, involves alterable representations after acquisition and displays experience-dependent characteristics, supporting that the genesis of AP can be understood from a learning perspective.

The learning perspective also provides a parsimonious account of the differential manifestations of AP ability for different individuals. Various subtypes of AP have been introduced to explain AP ability that is well above chance but not comparable with that of the so-called ‘genuine AP possessors’. For example, ‘pseudo AP’, ‘quasi-AP’, ‘implicit AP’, ‘latent AP’ and ‘residual AP’ have been used to refer to non-discarded AP information during development, unexpressed potential to develop AP limited by the critical period, incomplete forms of AP, etc. (Deutsch, 2013; Levitin & Rogers, 2005; Schellenberg & Trehub, 2003; Takeuchi & Hulse, 1993; W. D. Ward, 1999). These categorizations are established based on the assumption of a critical period in AP development. Given our challenge to the existence of the critical period on AP, it is doubtful how useful these categorizations are in enhancing our understanding of AP. Instead of being qualitatively different forms of AP, they may simply reflect different degrees and types of experience, and/or the attention and motivation in learning to process pitch information in an absolute manner. For instance, the ability to label the ‘A4’ tone only (referred as ‘quasi-AP’ or the ‘absolute A’) can be best explained by their habitual use of A4 as the tuning tone (Levitin & Rogers, 2005). The ‘imperfect’ or ‘incomplete’ AP performance may simply reflect various degree of AP learning, such as those demonstrated by the majority of the participants in this study (Figures 2 and 4). Research on whether there are qualitative or quantitative differences between different types of AP would be a fruitful future direction.

From the learning perspective, we should all have the potential to acquire AP regardless of age or prior musical training. While successful learning can be affected by many factors such as motivation and persistence (Vansteenkiste, Simons, Lens, Sheldon, & Deci, 2004), the difficulty to acquire AP (in terms of the slow improvement and relatively low success rate) is in clear contrast with the relative ease of other types of learning. For example, children can learn to name familiar objects in new languages fairly easily (Gathercole & Baddeley, 1990), and adults can learn to name tens of novel objects with nonwords in several hours (A. C.-N. Wong, Palmeri, & Gauthier, 2009; Y. K. Wong et al., 2011). What makes naming the twelve pitches so difficult to learn?

A possible reason concerns the interference from speech. Speech contains rich and fine-grained tonal information, such as pitch levels and pitch contours, that conveys different meanings and emotions (Saffran & Griepentrog, 2001). With years of experience in perceiving speech, tones and speech words may have formed intricate many-to-many mappings with a range of tones, or tonal patterns that are associated with different meanings and contexts. These experience-dependent mappings could be specific to each individual. It is possible that retrieving verbal labels of tones during a pitch-naming task activates the many-to-many mappings between tones and speech words, and that interferes with one’s judgment of the pitch name of the tone. This proposal is consistent with the observation that musicians performed worse with tones presented with human voice than other timbres (Vanzella & Schellenberg, 2010). The individual difference in speech interference could drive the individual variability in AP performance and acquisition. Future work can further explore this possibility in explaining pitch-naming performance in the general public.

## Conclusion

Overall, the current study shows that AP continues to be learnable in adulthood. The results challenge the influential theory that AP is only possible for few individuals with articular genes and training within the critical period. Instead the role of learning should be given a larger emphasis. The findings also form a good basis for future research on identifying, if any, aspects of AP more trainable in adulthood and aspects that are potentially exclusive for the few exceptional AP possessors.

## Acknowledgments

The authors declare no conflict of interest. We thank Gabriel Chan Pak Hong and Michael Lai Wei Chun for their help in data collection, Mandy Chu Yan Ting for the technical support, Helen Wong Hoi Shan for her help in violin tone production, and Patrick Bermudez for providing the synthetic tones.

## Authors Contributions

Y. Wong and A. Wong developed the study concept and designed the study. Y. Wong collected the data. Y. Wong, K. Lui and K. Yip analyzed the data. Y. Wong, K. Lui and A. Wong drafted the manuscript. All authors approved the final version of the manuscript for submission.

## Footnotes

1 A weaker definition of critical period refers to the early period of life during which experience has a particularly strong effect on development than the same experience at other times, while plasticity may still be observable and extend to adulthood (e.g., Hooks & Chen, 2007; Sengpiel, 2007). This definition makes the concept undifferentiable from ‘sensitive period’.

2 One participant did not participate in the testing one month later and was excluded from this analysis.

3 One musician and one non-musician did not participate in the testing three months later and were excluded from this analysis.

4 The performance pattern of this particular participant was highly inconsistent and uninterpretable. First, most participants tended to progress and improve more slowly as the training proceeded. However, this particular participant showed an opposite pattern. In particular, the progress of this participant was very slow in the first half of the training when only a few pitches were included. However, her progress suddenly and dramatically improved in the second half of the study when most of the pitches were included in the training. Also, she showed a large standard deviation in the number of attempts required to pass the training levels (*SD* = 55.4), which was more than two times higher than that of the rest of the group, showing that her performance was highly unstable and deviant of the learning patterns of others. Since the training was performed in an uncontrolled environment (unlike the previous experiments that were performed in the laboratory), the highly deviant learning pattern could be caused by many uninteresting reasons that are irrelevant to AP learning. Therefore, we decided to exclude the data of this participant for further data analyses.

5 One non-musician did not participate in the posttest one month later and was excluded from this analysis.

6 Four musicians were excluded from this analysis because they did not take part in any ABRSM before, therefore we did not have any data on their level of musical achievement.

7 We did not include the posttest at three months later because this test was not included in Experiment 1.

